# Large-scale metabarcoding analysis of epipelagic and mesopelagic copepods in the Pacific

**DOI:** 10.1101/2020.01.22.915082

**Authors:** Junya Hirai, Aiko Tachibana, Atsushi Tsuda

**Affiliations:** Atmosphere and Ocean Research Institute, The University of Tokyo, Chiba, Japan; Department of Ocean Sciences, Tokyo University of Marine Science and Technology, Tokyo, Japan

## Abstract

A clear insight into large-scale community structure of planktonic copepods is critical to understanding mechanisms controlling diversity and biogeography of marine taxa, owing to their high abundance, ubiquity, and sensitivity to environmental changes. Here, we applied a 28S metabarcoding approach to large-scale communities of epipelagic and mesopelagic copepods at 70 stations across the Pacific Ocean and three stations in the Arctic Ocean. Major patterns of community structure and diversity, influenced by water mass structures, agreed with results from previous morphology-based studies. However, large-scale metabarcoding approach could detected community changes even under stable environmental conditions, including changes in the north/south subtropical gyres and east/west areas within each subtropical gyre. There were strong effects of epipelagic environment on mesopelagic communities, and community subdivisions were observed in the environmentally-stable mesopelagic layer. In each sampling station, higher operational taxonomic unit (OTU) numbers and lower phylogenetic diversity were observed in the mesopelagic layer than in the epipelagic layer, indicating a recent rapid increase of species numbers in the mesopelagic layer. The phylogenetic analysis utilizing representative sequences of OTUs revealed trends of recent emergence of cold-water OTUs mainly distributed at high latitudes with low water temperatures. Conversely, high diversity of copepods at low latitudes was suggested to have been formed through long evolutionary history under high water temperature. The metabarcoding results suggest that evolutionary processes have strong impacts on current patterns of copepod diversity, and support the “out of the tropics” theory explaining latitudinal diversity gradients of copepods. Both diversity patterns in epipelagic and mesopelagic showed high correlations to sea surface temperature; thus, predicted global warming may have a significant impact on copepod diversity in both layers.

**Author Summary:** Marine planktonic copepods are highly dominant and diverse, and revealing their community structure and diversity is important to understanding marine ecosystems. We used molecular-based metabarcoding to reveal a total of 205 copepod communities in the ‘sunlight’ or epipelagic layer (0– 200 m) and the ‘twilight’ or mesopelagic layer (200–500 m and 500–1,000 m), mainly in the Pacific Ocean (data for 70 stations), but also in the Arctic Ocean (data for three stations). Different copepod communities were found in each geographical region with different environmental conditions, including tropical, subtropical, transition, Kuroshio Current, California Current, subarctic and arctic areas. The metabarcoding method sensitively detected small changes of copepod community even in environmentally-stable subtropical ocean systems and the mesopelagic layer. A high diversity of copepods was detected at low latitudes, and copepod diversity was higher in the mesopelagic layer than in the epipelagic layer in each area. These diversity patterns were influenced by both evolutionary history and present environmental conditions. The copepod community in the mesopelagic layer was strongly influenced by environmental conditions in the epipelagic layer. Thus, predicted climate changes may affect marine ecosystems not only in the epipelagic layer but also in the mesopelagic layer.

## Introduction

Zooplankton play a significant role as secondary or tertiary producers in pelagic communities, and their diversity is important in supporting the functions of marine ecosystems [1]. Approximately 7,000 species of marine zooplankton in 15 phyla have been described, and many additional undescribed species of zooplankton have been recently documented using molecular techniques [2, 3]. Marine planktonic copepods are an especially important group of zooplankton, having a high diversity with over 2,500 described species [4]. Their high abundance, ubiquitous distribution, and sensitivity to environmental changes also make copepods informative indicators in marine ecosystems [5]. Marine ecosystems are altered rapidly under climate change [6], and modifications in the biogeography of key copepod species have already affected higher-trophic levels, including commercially important fish [7]. Insights into the mechanisms that maintain copepod community structure and diversity would thus contribute to our understanding of broad-scale diversity patterns and monitoring changes in ocean ecosystems.

The Pacific Ocean, which covers more than 30 % of the Earth’s surface, is the largest ocean basin, providing numerous habitats for different marine organisms in complex ecosystems [8, 9]. As the Pacific Ocean was formed from the Panthalassic Ocean 160 million years ago (Ma), its basin therefore has a longer history than other ocean basins [10] and possibly harbors marine taxa with long evolutionary histories. In coastal marine ecosystems, for example, the western side of the Pacific facing the Coral Triangle is known as a hotspot of coastal marine species diversity [11]. A high diversity of oceanic copepods is thus expected in the Pacific; however, large-scale studies of copepod communities have mainly been restricted to the Atlantic Ocean [12–14]. A high diversity of copepods has been empirically estimated at low latitudes with high water temperatures and low productivity at a global scale; however, this estimation of diversity is primarily based on data from the Atlantic Ocean [15]. In early studies in the Pacific, only major species of zooplankton including copepods were examined for their distribution, and major water mass structures were a key factor in determining zooplankton biogeography [16, 17]. Whole-community analyses of copepods have only been carried out in specific areas, and there are relatively few large-scale studies in the Pacific; Williamson and McGowan compared copepod communities between north and south subtropical gyres [18], and Sun and Wang studied latitudinal patterns of copepods from the equator to subarctic regions along 160°E [19]. Calanoid copepod community and diversity have also been investigated from epipelagic (0–200 m) to bathypelagic (1,000–4,000 m) layers at 12 stations in the North Pacific [20]. However, large-scale copepod communities across the Pacific are especially limited below the epipelagic layer. Since pelagic ecosystems including copepod distributions are rapidly changing as a consequence of climate change [21], large-scale analyses of whole copepod communities with high taxonomic resolutions should be performed with high horizontal and vertical coverages in the Pacific Ocean and adjacent areas including the Arctic Ocean.

Large-scale studies of copepod diversity are limited because of difficulties in collecting samples over large areas. Previous large-scale studies of copepods have also required a high degree of expertise for morphological classifications, and not all taxa have been classified at the species-level. Molecular techniques with high taxonomic resolution are an effective tool for analyzing copepod diversity in the epipelagic and mesopelagic (200–1,000 m) layers, revealing cryptic and undescribed species in the open ocean [22, 23]. Metabarcoding, in particular, uses massive sequence data produced via high-throughput sequencers and is a promising technique to comprehensively describe zooplankton communities [24, 25]. This technique has already been applied to copepods using the Roche 454 high-throughput sequencer, revealing the broad-scale diversity and biogeography of copepods in the epipelagic layer of tropical and subtropical regions in the Pacific based on operational taxonomic units (OTUs) of 28S rRNA gene sequences [26, 27]. The metabarcoding method of copepods was further developed for use with the Illumina MiSeq platform, leading to low-cost and high-quality analyses of marine planktonic copepod diversity with high taxonomic resolutions [28]. In addition to high taxonomic resolution, metabarcoding analysis provides nucleotide sequences of OTUs, which are expected to provide insights into large-scale evolutionary processes and the reasons underlying patterns of copepod diversity. The global patterns of community and diversity have been analyzed using metabarcoding analysis of 18S rRNA gene for eukaryotic organisms including copepods [3, 29]; however, large-scale Pacific regions have not been fully covered, and detailed mechanisms controlling community structure and diversity should be discussed using the metabarcoding approach for both epipelagic and mesopelagic copepods in the Pacific Ocean.

In this study, we aimed to investigate the large-scale patterns of community and diversity of copepods based on genetic data using a metabarcoding approach in the Pacific Ocean as well as in the Arctic Ocean connected to the Pacific Ocean. Zooplankton samples were collected from the epipelagic layer (0–200 m) and the upper (200–500 m) and lower (500–1,000 m) mesopelagic layers from 2011 to 2017 at 70 stations across the Pacific Ocean as well as the epipelagic layer at three stations in the Arctic Ocean (Fig 1a; S1 Table). We performed metabarcoding using a high-throughput Illumina MiSeq sequencer on total 205 zooplankton samples, ranging from 40°S to 68°N and 40°E to 68°W covering Arctic, Subarctic, Transect, California Current, Kuroshio Current, North subtropical, South subtropical, Tropical regions. Broad-scale copepod communities were compared based on OTU data, revealing both vertical and horizontal patterns of copepod diversity. Subsequently, the environmental variables explaining copepod communities and diversity were evaluated. We also investigated the sequence diversity and phylogenetic relationships of the OTUs in order to reveal the effects of evolutionary processes on copepod diversity and biogeography in the Pacific Ocean.

**Fig 1.**
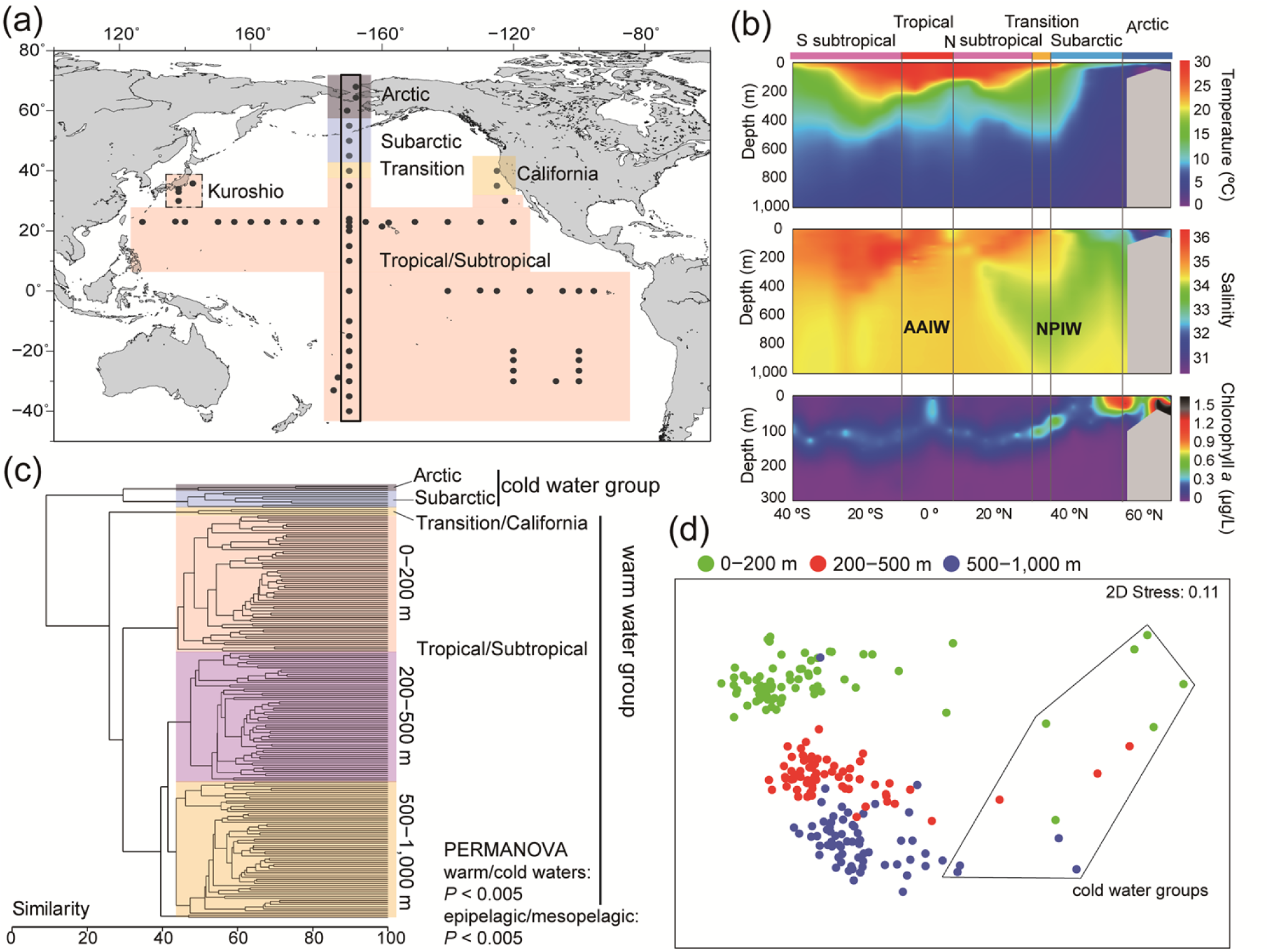
Large-scale community compositions of copepods based on the presence/absence of operational taxonomic units (OTUs) in the Pacific and Arctic Oceans. (a) Sampling location. (b) Vertical profiles of environmental variables across latitudes (top: seawater temperature, middle: salinity, and bottom: chlorophyll-*a*) obtained during north–south transect cruises along 170°W, which is surrounded by black lines in (a); the North Pacific Intermediate Water (NPIW) and Antarctic Intermediate Water (AAIW) are shown in the salinity profile. (c) Cluster analysis of all copepod communities. The boundaries of warm-, transition/California-, and cold-water communities in the epipelagic layer are illustrated in (a). (d) Multidimensional scaling analysis of all samples. Cold-water communities are surrounded by black lines. The PERMANOVA was performed for copepod community compositions between cold and warm waters and between epipelagic and mesopelagic layers. Details of the PERMANOVA are presented in S3 Table.

## Results

### Metabarcoding data

In this study, we used a criterion of OTUs at the 98.5 % similarity threshold and a minimum sequence reads for OTUs of ≥ 8, to avoid overestimating diversity and to maintain an appropriate taxonomic resolution. These values of similarity and abundance thresholds were determined based on the preliminary analyses using mock communities (S2 Table). Mock communities contained morphologically-identified copepod species, and OTUs at different similarity and abundance thresholds were compared with reference sequences of copepod species obtained by Sanger sequencing. The mock community data samples were also analyzed together with all sequence data of environmental communities for validating accuracy of bioinformatics (S1 Fig). A total of 14,117 sequence reads were obtained from each sample after a standardized number of sequence reads in the analyses of environmental communities, and 2,893,985 copepod sequence reads were clustered into 1,659 copepod OTUs. The rarefaction curve using all sequence data showed that numbers of OTUs reached a plateau (Fig 2a). Not all rarefaction curves of each sample at each sampling layer (epipelagic; upper mesopelagic; lower mesopelagic) reached a plateau in numbers of OTUs especially at low latitudes (e.g., Tropical and Subtropical) or in the mesopelagic layer; however, spatial patterns of diversity, including differences of geographical areas and sampling layers, were reflected in numbers of OTUs in our datasets (Fig 2b).

**Fig 2.**
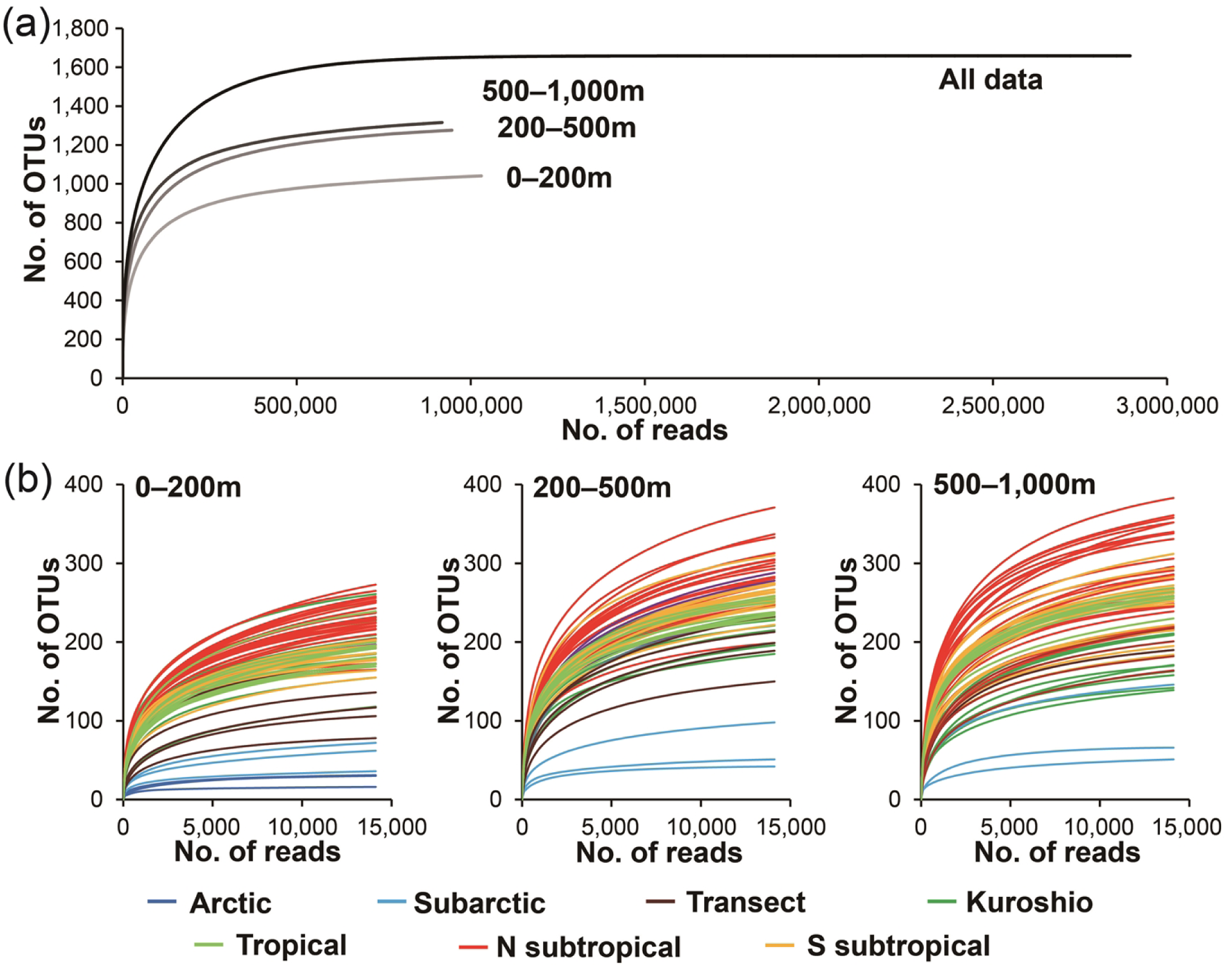
Rarefaction curves for OTU numbers at different sequence reads. (a) Results for all sequence reads and total sequence reads at each sampling layer (0–200 m, 200–500 m, and 500– 1,000 m. (b) Results for each sampling layer. Different geological areas are represented by different color.

### Copepod compositions based on presence/absence of OTUs

Water temperature, salinity, and chl-*a* concentrations varied among the warm waters at low latitudes and cold waters in the high latitudes, especially in the epipelagic layer (Fig 1b). These environmental changes led to significantly different copepod compositions based on presence/absence of OTUs between warm and cold waters at each sampling layer (*P* < 0.005; Fig 1c; S3 Table). In the cold-water group, compositions in the arctic and subarctic were clustered into different groups. Copepod compositions were clearly differentiated by sampling depth between the epipelagic and mesopelagic layers (*P* < 0.005; Fig 1d). The transition zone and the California Current region in the epipelagic layer formed a distinct group, separated from other warm-water groups, including Tropical, North and South subtropical, and Kuroshio regions.

The clustered groups of copepod compositions in the warm-water group clearly corresponded to the respective geographical areas in each sampling layer, including transition, California Current, Kuroshio, subtropical, and tropical regions (Fig 3). The latitudinal subdivisions of community compositions were especially evident in the epipelagic layer even within subtropical regions (Fig 3a; *P* < 0.005), including differences between the North and South Pacific subtropical regions. These community changes were reflected in the best model, which showed that the average water temperature in the epipelagic layer was the factor most strongly related to epipelagic copepod compositions, followed by latitude (Table 1). The other geographical factor of longitude was also included in the best model for the epipelagic layer.

**Fig 3.**
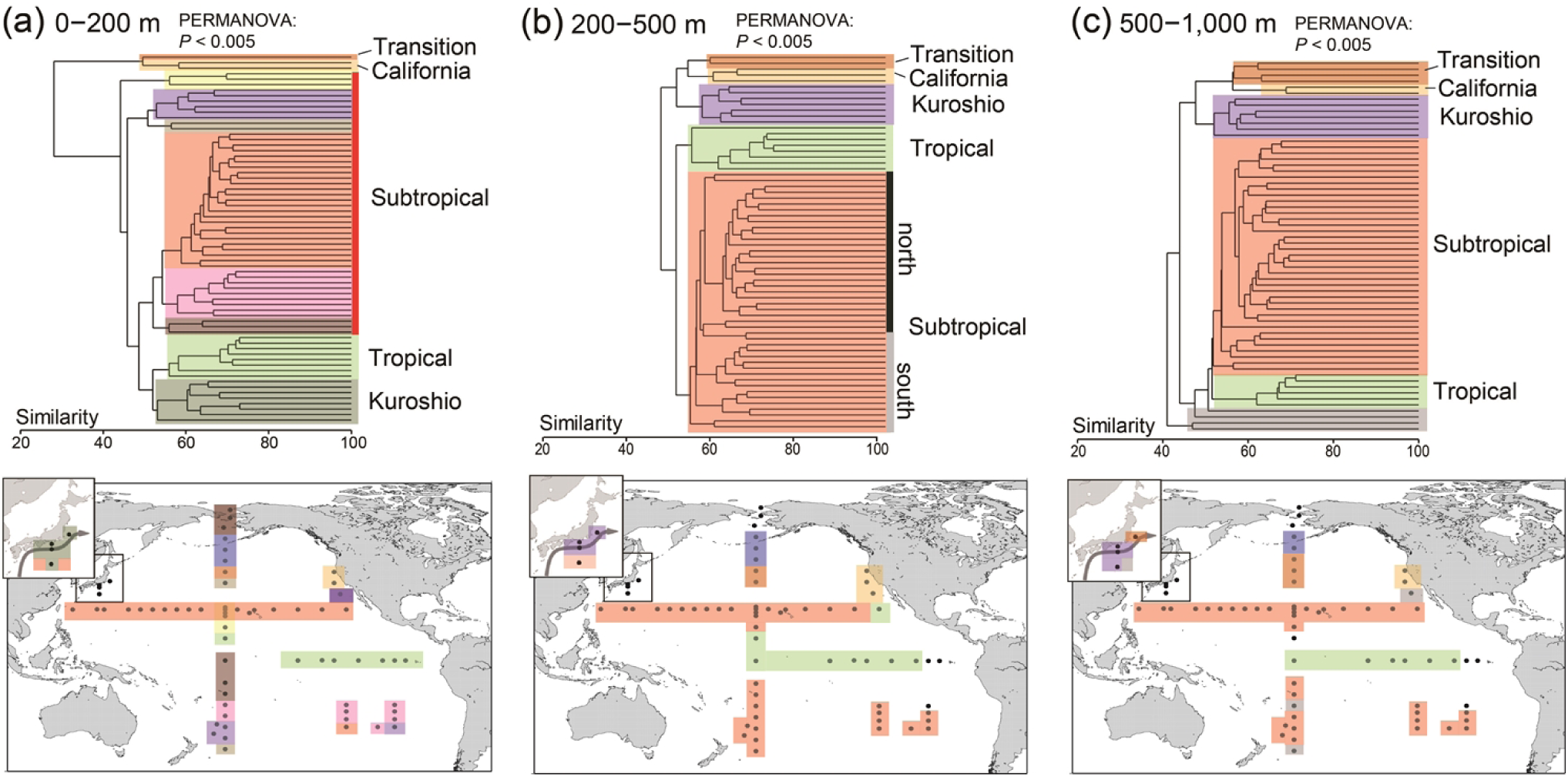
Community compositions of copepods at each layer based on presence/absence of operational taxonomic units (OTUs). (a) Epipelagic layer (0–200 m). (b) Upper mesopelagic layer (200–500 m). (c) Lower mesopelagic layer (500–1,000 m). Cluster analyses were performed on warm-water groups. The PERMANOVA was performed on groups represented by different colors at each sampling layer, except for the unclassified groups (gray) in the lower mesopelagic layer. Details of the PERMANOVA are presented in S3 Table.

**Table 1.** Summary of distance-based linear model (DistLM) analyses for copepod communities based on presence/absence of OTUs. The best model of variables explaining the copepod compositions was selected based on Akaike information criteria (AICs) for all locations in each sampling layer [epipelagic (0–200 m), upper mesopelagic (200–500 m), lower mesopelagic (500– 1,000 m), and throughout the sampling layers (0–1,000 m)]. The pseudo–*F*, *P*–value, and explained variation attributable to the model are indicated for each variable.

| | Variable | Pseudo-$F$ | $P$-value | Variation (%) |
| --- | --- | --- | --- | --- |
| **0–200 m** | Temp. (0–200 m) | 14.0 | 0.001 | 16.9 |
| AICc: 525.0 | Latitude | 9.9 | 0.001 | 10.5 |
| Variations: 35.5% | DO (0–200 m) | 5.7 | 0.001 | 5.7 |
|  | Longitude | 2.5 | 0.006 | 2.4 |
| **200–500 m** | Salinity (0–200 m) | 7.8 | 0.001 | 11.1 |
| AICc: 470.8 | Temp. (0–200 m) | 7.9 | 0.001 | 10.0 |
| Variations: 28.8% | DO (0–200 m) | 4.0 | 0.001 | 4.8 |
|  | Longitude | 2.4 | 0.004 | 2.9 |
| **500–1,000 m** | Latitude | 7.0 | 0.001 | 10.3 |
| AICc: 462.0 | Temp. (0–200 m) | 5.5 | 0.001 | 7.6 |
| Variations: 27.5% | DO (0–200 m) | 4.1 | 0.001 | 5.3 |
|  | DO (500–1,000 m) | 3.4 | 0.001 | 4.3 |
| **0–1,000 m** | Temp. (0–200 m) | 16.3 | 0.001 | 21.0 |
| AICc: 424.8 | Longitude | 6.6 | 0.001 | 7.9 |
| Variations: 38.9% | DO (0–200 m) | 4.9 | 0.001 | 5.5 |
|  | DO (0–1,000 m) | 4.3 | 0.001 | 4.5 |

Temp = average temperature; DO = average dissolved oxygen.

Although variations of community compositions in the upper and lower mesopelagic layers were relatively small compared with those of the epipelagic layer, we also observed different groups clustered by geographical regions in the upper and lower mesopelagic layers (*P* < 0.005; Fig 3b–c). The differences between the North and South Pacific subtropical regions were clearer in the upper mesopelagic layer than in the lower mesopelagic layer. The composition in the Kuroshio region showed high similarity to those in transition and California Current regions in the mesopelagic layer, regardless of the similar structures between Kuroshio and tropical regions in the epipelagic layer. The epipelagic environment had a strong effect on the compositions in the mesopelagic layer (Table 1). The compositions in the upper mesopelagic layer were explained by average salinity, water temperature, and dissolved oxygen content of the epipelagic layer as well as longitude in the best model. In the lower mesopelagic layer, latitude was the variable explaining most of the variation in composition, followed by average water temperature, dissolved oxygen content in the epipelagic layer, as well as the dissolved oxygen content in the lower mesopelagic layer.

The groups based on community compositions for the epipelagic and mesopelagic layers together (0–1,000 m) showed clear geographic changes (Fig 4a–c; *P* < 0.005). In the Kuroshio region, the inshore region of the Kuroshio Current was discriminated from the oceanic region. Community compositions corresponding with geographical changes were observed within the subtropical regions, including differences between the North and South Pacific. In addition to latitudinal changes, the western and eastern sides of subtropical gyres were clustered into different groups within North and South Pacific subtropical gyres. These clustered groups corresponded with distributions of water mass structures in the Pacific, as represented by the profiles of T-S diagrams at 0–1,000 m (Fig 4d). The latitudinal and longitudinal changes through water column were also observed in the best model to explain changes of copepod composition. The variable explaining most of the variation in community compositions at 0–1,000 m was average water temperature in the epipelagic layer, followed by longitude. Among cluster groups through the water column, there is a clear difference between warm and cold waters in the taxonomic compositions in each layer (Fig 4a). There are no clear differences with respect to family-level taxonomic compositions within warm waters, and proportions of unclassified copepods were high in the mesopelagic layer.

**Fig 4.**
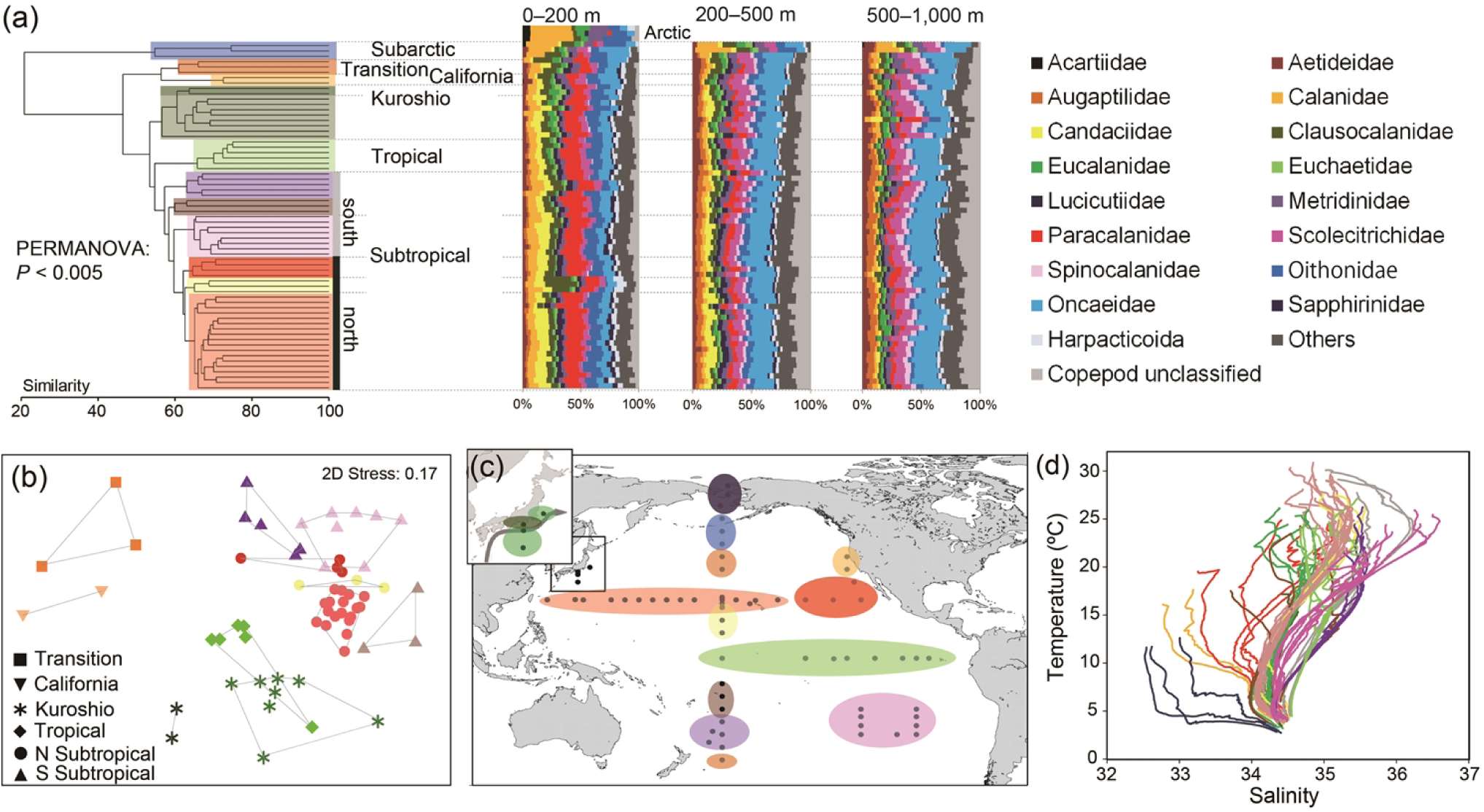
Community compositions of copepods based on presence/absence of operational taxonomic units (OTUs) throughout the sampling layers (0–1,000 m). (a) Cluster analysis. The PERMANOVA was performed on the groups represented by different colors. Taxonomic compositions of OTUs at each layer are shown for each sampling station in cluster analysis. The arctic community is also added to the epipelagic layer. Details of the PERMANOVA are presented in S3 Table. (b) Multidimensional scaling analysis for warm-water groups. Cluster groups are surrounded by lines. (c) Clustered groups visualized on map. (d) Comparison of clustered groups and T-S diagrams to investigate effect of water mass structures on copepod compositions.

### Copepod community based on sequence reads

The distribution peaks of major OTUs largely affected large-scale copepod community based on quantitative data using relative proportions of sequence reads. In addition to compositions based on presence/absence of OTUs, overall cluster analysis based on sequence reads also showed significantly different groups, which corresponded with sampling depth and geographical regions (*P* < 0.005; Fig 5). The key variables that explained copepod communities based on sequence reads were similar to those explaining the presence/absence of OTUs; in particular, environmental conditions in the epipelagic layer affected copepod communities based on sequence reads both in the epipelagic and mesopelagic layers (S4 Table). A total of 36 OTUs were selected as major OTUs to contribute to differences among cluster groups (Fig 5). These major OTUs showed a distinct difference between cold and warm waters, and a small number of specific OTUs dominated the arctic and subarctic regions in the cold-water group as well as in the transition-California region. These OTUs were mainly restricted to high latitudes, although some OTUs were present in warm-water mesopelagic layers especially in the Kuroshio region (e.g., OTU 4 with 100% identity to *Oithona similis*). Warm-water OTUs were widely distributed in each layer at low latitudes; however, peaks of sequence reads were different in each OTU even in closely-related taxa. For example, in the family Paracalanidae, which was dominant in the warm-water epipelagic layer, OTU 7 (100% identity to *Delibus nudus*) and OTU 19 (100% identity to *Paracalanus aculeatus*) showed high proportions of sequence reads in subtropical and tropical regions, respectively. Different distribution patterns were also observed in the family Scolecithrichidae in the mesopelagic layer, including OTU 36 (97% identity to *Scaphocalanus similis*) and OTU 68 (100% identity to *S. magnus*). We also detected specific OTUs with high dominance mainly in the Kuroshio region, including the genera *Subeucalanus* (OTUs 43, 115, and 144) and *Rhincalanus* (OTUs 12 and 52).

**Fig 5.**
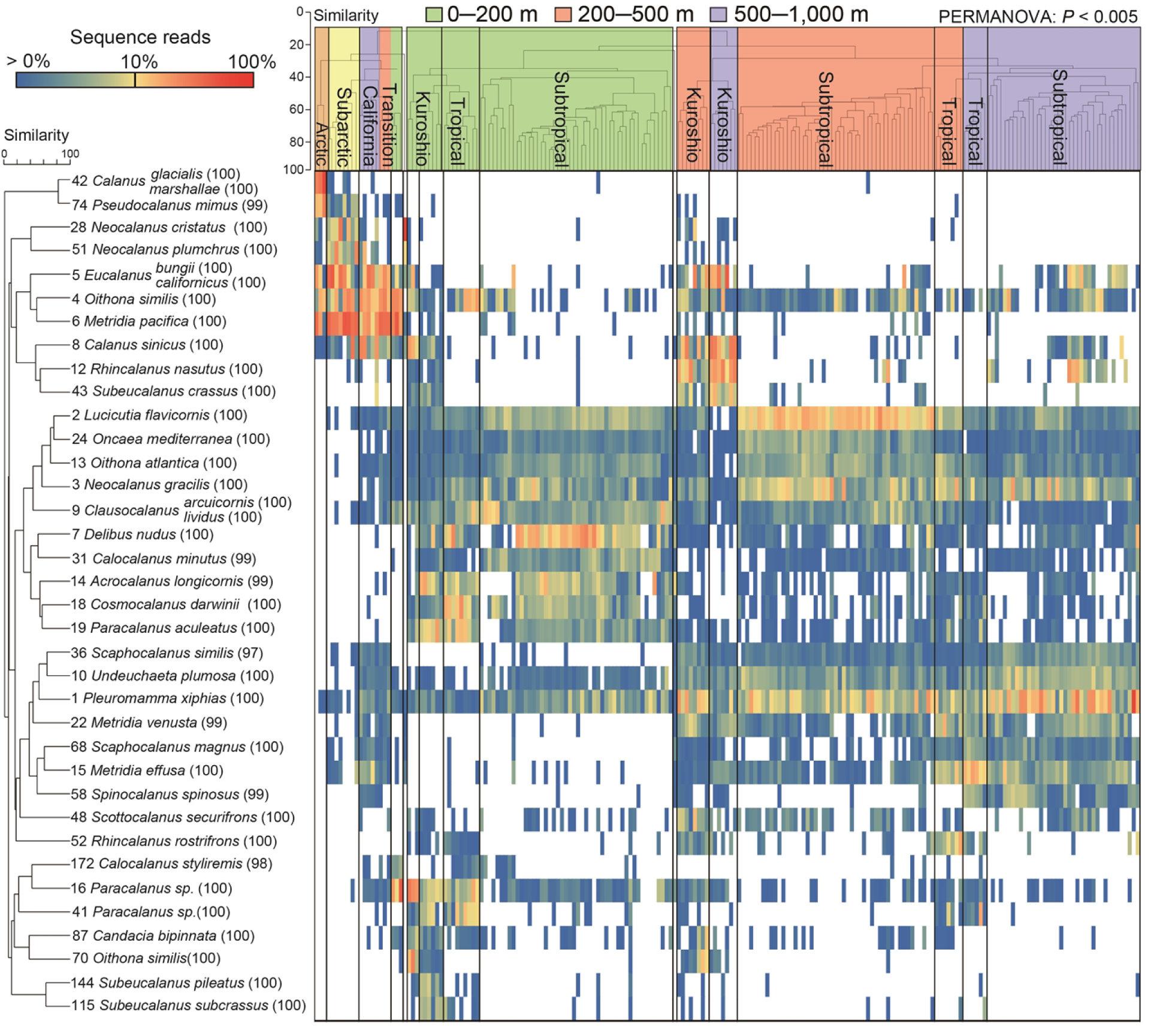
Distribution of quantitative data of sequence reads in major operational taxonomic units (OTUs). The cluster analysis was performed on all samples based on proportions of sequence reads in OTUs, and the PERMANOVA was performed on clustered groups. The OTU numbers and the best-hit species resulting from the BLAST search are represented for major OTUs, which were selected by the SIMPER analysis. The numbers in parentheses indicate the similarity percentage of the best-hit species. The proportions of sequence reads are log-transformed.

### Spatial patterns of the copepod diversity

OTU numbers in each layer showed clear horizontal changes, with larger OTU numbers at warm-water low latitudes than at cold-water high latitudes (Fig 6). There was an asymmetric pattern of latitudinal gradients in OTU numbers between the Northern and Southern hemispheres, and significantly higher (*P* < 0.05) OTU numbers were observed in the subtropical North Pacific than in other regions (Fig 7a; S5 Table). There was no clear difference in OTU number between the South subtropical and tropical regions. The Kuroshio region temporarily showed large OTU numbers in the epipelagic layer; however, low diversity was observed in the lower mesopelagic layer in the Kuroshio region compared with tropical and subtropical regions. These spatial patterns of diversity were also observed in the rarefaction curves for each layer (Fig 2). The variable explaining most of the variation in the spatial patterns of OTU numbers in all layers was sea surface temperature (SST) (Table 2). For the epipelagic layer, after SST, salinity, mixed layer depth (MLD), chlorophyll-*a* (chl-*a*), dissolved oxygen content, and latitude were selected in the best model. For the mesopelagic layer, after SST, longitude was the variable that explained most of the variation. In addition to SST, environmental factors of the epipelagic layers were also included in the best models for the mesopelagic layer and throughout the sampling layers, including MLD, dissolved oxygen content, and chl-*a*.

**Fig 6.**
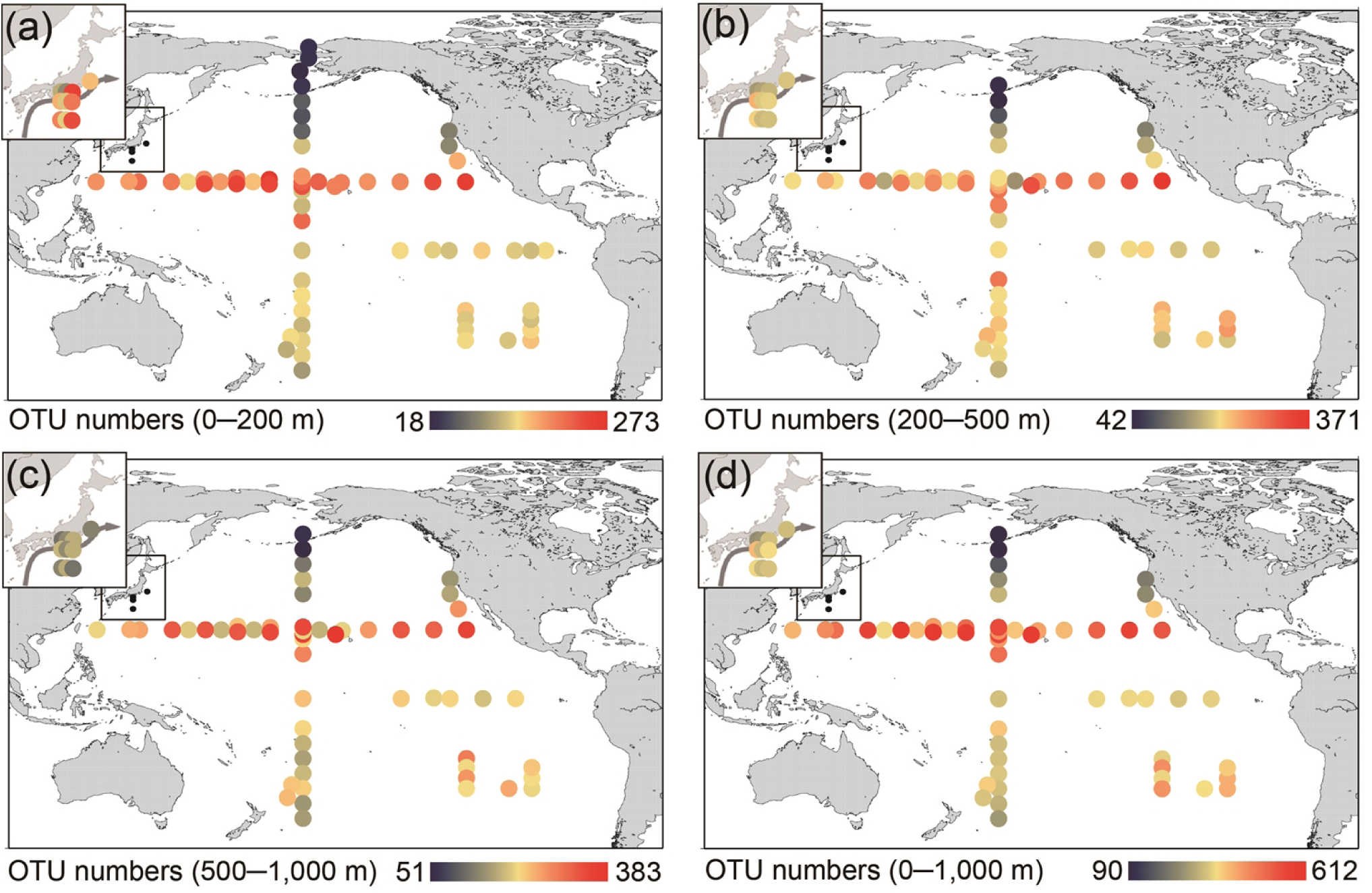
The distribution of copepod diversity in the Pacific Ocean at different layers of the water column: (a) epipelagic (0–200 m), (b) upper mesopelagic (200–500 m), (c) lower pelagic (500– 1,000 m), and (d) all sampling layers (0–1,000 m). Colors indicate the number of operational taxonomic units (OTUs).

**Fig 7.**
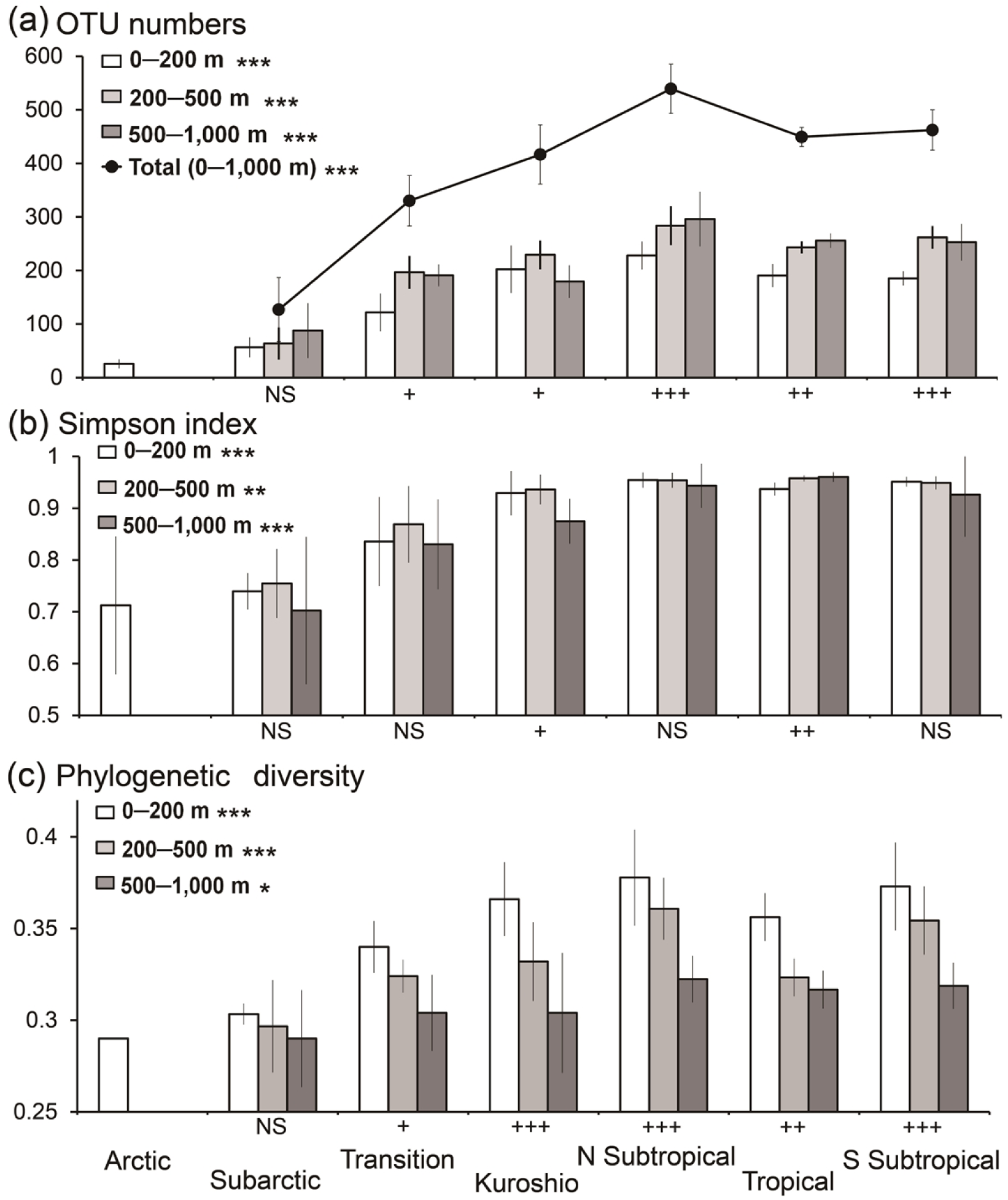
Horizontal and vertical patterns of copepod diversity. (a) Operational taxonomic units (OTUs). Total OTUs were calculated for each sampling location in all layers (0–1,000 m). (b) Simpson diversity index. (c) Phylogenetic diversity. The California Current region is included in Transition. The diversity parameters were averaged for each region and scale bars indicate SD. NS = not significant; * = *P* < 0.05; ** = *P* < 0.005; *** = *P* < 0.001 based on Kruskal-Wallis tests for comparisons among geographic regions. NS = not significant; + = *P* < 0.05; ++ = *P* < 0.005; +++ = *P* < 0.001 for Kruskal–Wallis tests for comparisons among sampling depths. Details of statistical analyses are shown in S5 Table.

**Table 2.** Variables explaining operational taxonomic unit (OTU) numbers. The best model of variables was selected using a stepwise method in generalized linear model (GLM) analysis, and the pseudo R^2^ and Akaike information criteria (AIC) values are reported for the best models.

|  | Variable | Estimate | Z value | P |
| --- | --- | --- | --- | --- |
| **0–200 m** | Intercept | 1.53 | –0.69 | NS |
| Pseudo R<sup>2</sup>: 0.72 | SST | 4.28E–02 | 5.21 | < 0.001 |
| AIC: 735.42 | Salinity (0–200 m) | 1.69E–01 | 2.54 | < 0.05 |
|  | MLD | 2.38E–03 | 2.41 | < 0.05 |
|  | Chl-*a* | –1.28E–01 | –2.03 | < 0.05 |
|  | DO (0–200 m) | –6.11E–02 | –1.85 | NS |
|  | Latitude | 2.40E–03 | 1.44 | NS |
| **200–500 m** | Intercept | 3.35 | 11.39 | < 0.001 |
| Pseudo R<sup>2</sup>: 0.58 | SST | 4.49E–02 | 6.74 | < 0.001 |
| AIC: 693.35 | Longitude | 2.49E–03 | 3.38 | < 0.001 |
|  | MLD | 2.20E–03 | 2.48 | < 0.05 |
|  | DO (0–200 m) | 6.86E–02 | 1.92 | NS |
|  | Temp (200–500 m) | 1.66E–02 | 1.64 | NS |
| **500–1,000 m** | Intercept | 3.48 | 14.50 | < 0.001 |
| Pseudo R<sup>2</sup>: 0.65 | SST | 4.57E–02 | 9.56 | < 0.001 |
| AIC: 617.34 | Longitude | 2.80E–03 | 5.88 | < 0.001 |
|  | DO (500–1,000 m) | –5.46E–02 | –4.05 | < 0.001 |
|  | DO (0–200 m) | 1.11E–01 | 3.53 | < 0.001 |
| **0–1,000 m** | Intercept | 10.08 | 2.76 | < 0.01 |
| Pseudo R<sup>2</sup>: 0.77 | SST | 4.49E–02 | 9.68 | < 0.001 |
| AIC: 626.71 | Longitude | 2.52E–03 | 4.75 | < 0.001 |
|  | DO (0–200 m) | 1.51E–01 | 4.18 | < 0.001 |
|  | Temp (0–1,000 m) | 3.04E–02 | 3.01 | < 0.01 |
|  | Chl-*a* | 6.39E–01 | 2.81 | < 0.01 |
|  | MLD | 1.07E–03 | 2.48 | < 0.05 |
|  | Salinity (0–1,000 m) | –1.89E–01 | –1.74 | NS |
|  | DO (0–1,000 m) | –3.63E–02 | –1.62 | NS |

SST = sea surface temperature, Temp = average water temperature (°C), Chl-*a* = average chlorophyll-*a* concentrations in the epipelagic layer (0–200 m) (µg/L), DO = average dissolved oxygen (ml/L), MLD = mixed layer depth (m), NS = not significant.

There was a regional pattern of high diversity at low latitudes for the Simpson diversity index (Fig 7b) and phylogenetic diversity (Fig 7c). Unlike the OTU numbers, the Simpson index values showed an unclear north–south asymmetric pattern for each layer in the tropical and subtropical regions. Phylogenetic diversity, which were average sequence variations among OTUs based on genetic distance, was high in the North and South Pacific subtropical regions for each layer. The regional differences of phylogenetic diversity were especially evident in the epipelagic and upper mesopelagic layers, and significantly high phylogenetic diversity (*P* < 0.05) was observed especially in the North subtropical region compared with other regions (S5 Table).

In addition to horizontal patterns, a vertical gradient of copepod diversity was observed, with significantly higher OTU numbers in the mesopelagic layers than in the epipelagic layer in most geographic regions (Fig 7a; S5 Table). No clear differences of OTU numbers were found between the upper and lower mesopelagic regions in each geographic region. A vertical gradient of diversity was not evident in the Simpson diversity index (Fig 7b). However, there was a clear vertical gradient in phylogenetic diversity, with decreasing phylogenetic diversity with increasing depth in each geographic region (Fig 7c). The phylogenetic diversity was significantly higher (*P* < 0.05) in the epipelagic layer than in lower mesopelagic layers in all regions except the subarctic (S5 Table). Different spatial and vertical distribution patterns of OTUs were associated with taxonomic groups of copepods (Fig 8). High proportions of OTU numbers with distributions in the mesopelagic layer were found in taxonomically diverse groups such as the superfamilies Augaptiloidea, Bathpontioidea, and Spinocalanoidea, as well as in the families Megacalanidae, Rhincalanidae, Aetideidae, and Scolecitrichidae in calanoid copepods. Non-calanoid copepods including those belonging to the families Lubbockiidae and Oncaeidae in Cyclopoida, and those belonging to the order Mormonilloida also had high proportions of OTUs with the main distribution in mesopelagic layers. These taxonomic groups essentially showed large OTU numbers, and these taxonomic groups were also abundant with high proportions of OTU compositions in the mesopelagic layer (Fig 4a). Taxonomic families with high proportions of OTUs distributed in the epipelagic layer included Centropagedae, Pontellidae, Temoridae, Paracalanidae, and Clausocalanidae in the order Calanoida, and Corycaeidae in the order Cyclopoida.

**Fig 8.**
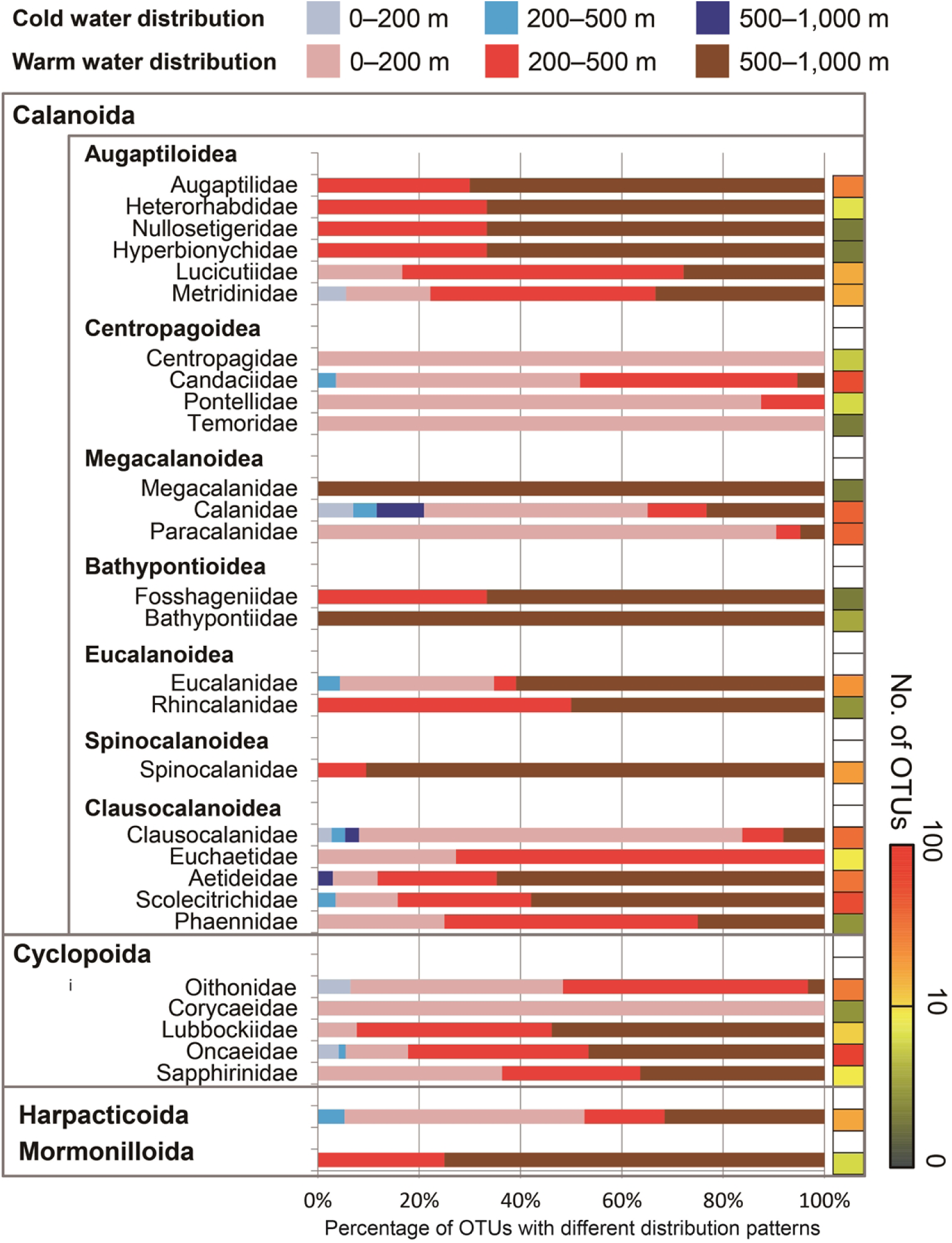
Distribution patterns of copepod OTUs based on cold- and warm-water groups and water depth [epipelagic (0–200 m), upper mesopelagic (200–500 m), and lower mesopelagic (500–1,000 m)]. Only operational taxonomic units (OTUs) with ≥ 15 sequence reads in a single sample were analyzed, and proportions of numbers of OTUs with different distribution patterns are shown in each taxonomic group. The logged-scale total OTU numbers are represented as colors for each taxonomic group. The order Calanoida and Cyclopoida were further classified into taxonomic families. The superfamily names are indicated for the order Calanoida.

OTUs with distribution peaks in cold waters at high latitudes including arctic and subarctic regions were detected in different taxonomic groups of copepods, which were further used for phylogenetic analyses to investigate effects of phylogenetic relationships on OTU distribution patterns (Fig 9). In the phylogenetic tree of taxa containing both cold-water and warm-water OTUs, bootstrap values were relatively low in most groups; however, basal positions were mostly occupied by warm-water OTUs with large OTU numbers rather than by cold-water OTUs. The average nearest genetic distances, which could be used as a proxy of divergent time between closely-related OTUs, were lower for cold-water OTUs than for warm-water OTUs, except for Clausocalanidae and Oncaeidae. Similar genetic distance values were observed in the cold-water and warm-water OTUs of Eucalanidae.

**Fig 9.**
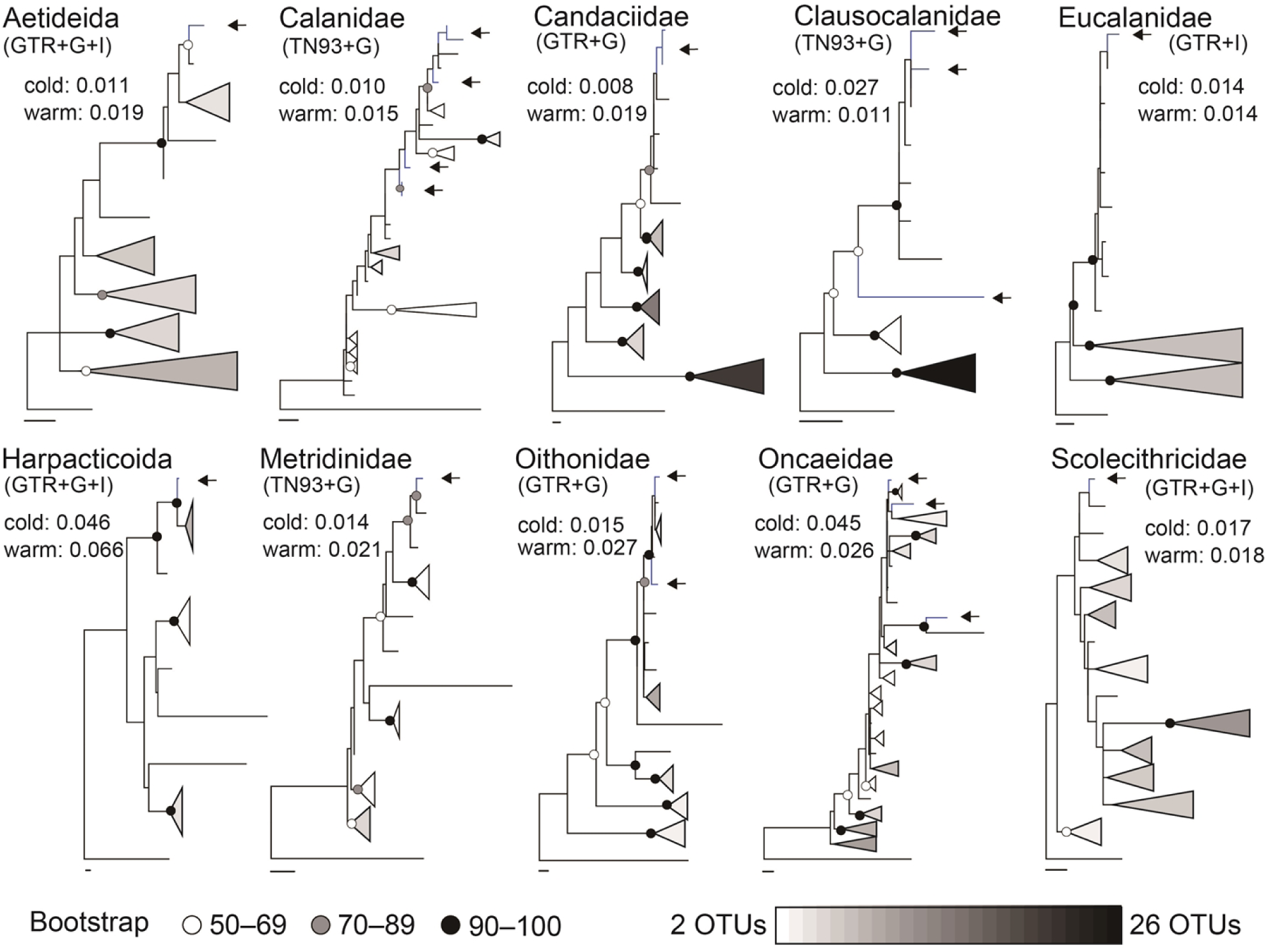
Phylogenetic analysis of taxonomic groups including operational taxonomic units (OTUs) with warm-water and cold-water distributions. OTUs with cold-water distributions are represented by arrows. Nodes show bootstrap values from maximum likelihood analysis (white, grey, black). Triangles are genetic lineages containing multiple OTUs with warm-water distributions to simplify the phylogenetic trees, and OTU numbers within each triangle are represented in the white-black spectrum. The length of triangle varies according to the deepest node of each lineage. Scale bars indicate 0.02 genetic distances for each tree. The average nearest genetic distance in OTUs is indicated for copepods with warm-water and cold-water distributions in each taxonomic group. The nucleotide substitution models used are indicated for each taxon (GTR: General Time Reversible; TN93: Tamura-Nei).

## Discussion

To the best of our knowledge, this is the first study to present large-scale distribution patterns of copepods focusing on both epipelagic and mesopelagic layers of the Pacific Ocean using massive sequence data. Additionally, no previous metabarcoding studies of zooplankton have combined distribution patterns with comprehensive sequence data to explain evolutionary processes. We selected the 28S D2 region in order to obtain taxonomic resolutions higher than 18S rRNA gene sequences, and higher primer specificity than mitochondrial genes (e.g., cytochrome oxidase I), which are common for DNA barcoding of zooplankton [2]. The 28S rRNA gene is also useful for studies of the phylogenetic relationship, with high of database coverage for various taxonomic families [30], and metabarcoding analysis using the 28S rRNA gene can be used to detect a wide range of taxonomic groups of copepods using primer pairs [27]. In this study, the utility of metabarcoding method was validated using mock community analyses; the rarefaction curve using all sequence data reached a plateau in our study. A metabarcoding technique is shown to be a powerful tool to study community structure and diversity of copepods; however, there are methodological limitations in metabarcoding, such as overestimation of OTUs and quantitative biases [24]. In addition to adding a reference library of DNA barcoding especially in mesopelagic copepods, future developments of sequencing technology and bioinformatic methods are necessary to improve metabarcoding of zooplankton. Although there are limitations in the current metabarcoding analysis of copepods, the extensive data presented in this study is valuable as the first large-scale study providing an overview of copepod biogeography and diversity from a genetic aspect for the Pacific and Arctic Oceans.

In previous studies using morphological classification, a total of 125 calanoid copepod species were reported from 0–600 m in the North Pacific gyre [31], and 211 calanoid species were reported from 0–2,615 m in the tropical to subarctic regions of the North Pacific [20]. Although numbers of OTUs and morphological species cannot be compared directly, the patterns of OTUs obtained in this study are good proxies of species diversity. A total of 1,659 OTUs from approximately three million reads thus indicates the hitherto hidden diversity of copepods. Previously, over 5,000 copepod OTUs have been reported in the epipelagic layer of the global oceans [3], and 1,806 copepod OTUs have been reported from 0–1,500 m in the North Pacific subtropical gyre [32]. Differences in total OTU numbers are possibly related to different sampling and experimental methods, bioinformatic strategies, and spatial coverage of samples. It should be noted that we mainly used a standardized method for sampling using the same sampling gear, with some exceptions in the Arctic and Kuroshio area (S1 Table). Although the mesh size of the plankton net, which largely affect results of metabarcoding analysis of zooplankton [33], were same in all stations, different size mouth openings for the sampling gear may have affected regional differences of copepod communities in this study.

The major groups, based on both quantitative (sequence reads) and non-quantitative (presence/absence of OTUs) analyses, included the distinct communities in the major water masses of arctic, subarctic, transition, subtropical, and tropical regions; the results are consistent with the results of previous studies using morphological classifications [12, 20, 31, 32]. The dominance and compositions of copepods are thus strongly influenced by geological areas with different environmental conditions. The western boundary of the Kuroshio and California Currents led to similar community structures in the equatorial and transition areas in the epipelagic layer, respectively, because of the ocean current systems in the Pacific. In addition to major community patterns, our molecular-based approach revealed subdivisions of copepod compositions even in the environmentally-stable areas, including the east and west subdivisions within subtropical gyres. Although there are a lot of common morphological species between the North and South Pacific subtropical gyres [18], copepod compositions in North and South subtropical areas were differentiated in our study, and latitudinal subdivisions of community structures were observed within each subtropical gyre. These geographically separated communities were supported by the best model to explain copepod community compositions including both environmental (e.g., temperature) and geographic factors (e.g., latitude and longitude). Previous studies have shown that gene flow of copepods is limited even in oceanic areas [34] and whole planktonic communities are strongly influenced by distances associated with ocean currents [35]. Although warm-water species are widely distributed at low latitudes, in this study there were species-specific distribution peaks in dominant OTUs. Zooplankton adaptive evolution may occur faster than previously considered [36]. The copepod community, thus, might have been subdivided based on our molecular-based data through adaptation to geographically separated local environments with different water masses in the Pacific and Arctic Oceans.

As well as subdivisions of copepod communities within subtropical areas, there were large-scale patterns of community structures found in the mesopelagic layer with relatively stable environments. The changes in copepod community composition were especially evident in the epipelagic layer, where environmental changes were large. These epipelagic environmental changes were also shown to affect community structures and the diversity of copepods in the mesopelagic layer. In addition to influences of particle flux from the epipelagic layer [20, 37], there are mesopelagic copepods with diel vertical migration to the epipelagic layer. Some copepods also migrate to the mesopelagic layer in order to avoid unfavorable environmental conditions in the epipelagic layer even in the warm waters, including those in the Kuroshio region [38]. A significant change of biomass and trophic efficiency is predicted to occur in mesopelagic nekton as well as in large zooplankton as a result of climate change [39]; thus, interactions between the epipelagic and mesopelagic layers are key regulators of local communities and diversity patterns in mesopelagic copepods.

Pelagic fauna typically display a latitudinal diversity gradient, with high species numbers in warm low-latitude waters and declining diversity toward higher latitudes [40]. Although a previous study of prokaryotes and eukaryotes using metabarcoding and image data showed no latitudinal diversity in the mesopelagic layer [29], our metabarcoding approach revealed clear latitudinal gradients of diversity in each layer and throughout the sampling layers. As shown by the diversity index and proportions of sequence reads, many phylogenetically-diverse species coexisted in oligotrophic warm waters at low latitudes, whereas fewer species dominated in the cold, food-rich waters at high latitudes, consistent with the estimates of global diversity of epipelagic copepods based on the morphological classifications [15, 41]. In spite of similar environmental conditions in the north and south subtropical gyres, we detected north–south asymmetry in OTU numbers, with a peak observed in the north subtropical region. In addition to the complex water mass structures in the large area of the North Pacific, including a strong western boundary current and the interaction of two major intermediate waters, the inside North Pacific subtropical gyre has been known as a place where floating materials collect [42, 43]. Therefore, the co-existence of more diversified copepods in complex ecosystems might contribute to higher species richness in the North Pacific than in the South Pacific.

Sequence data combined with OTU distributions provided insights on mechanisms forming the large-scale patterns of copepod diversity. Among the several hypotheses that have been proposed to explain latitudinal diversity gradients, our study is mainly consistent with the “out of the tropics” model, in which warm tropical regions are characterized by high speciation and low extinction rates, and most extra-tropical species belong to lineages that originated in the tropics [44]. While our molecular data cannot provide insight on extinction, our phylogenetic analyses and nearest genetic distance data support the hypothesis by van der Spoel and Heyman [42] that the central water (subtropical gyre) species first emerged in planktonic taxa ancestral to cold-water fauna. Most of the current planktonic zooplankton are hypothesized to have evolved and assembled after the K-T boundary mass extinction ∼65 Ma [45], when the earth’s temperature steadily decreased and latitudinal environmental gradients became stronger [46]. Although an appropriate molecular clock is not available for planktonic copepods, high-latitude species of each taxonomic group are likely to have evolved through adaptation to latitudinal environment gradients, such that low diversity at high latitudes may reflect this recent emergence of cold-water groups. The high metabolic rate and short generation time of copepods at high temperatures [1] also might have enabled rapid molecular evolution and high diversity at low latitudes over a long evolutionary history in the historically stable subtropical gyres [47]. Because this study only used short fragments of 28S sequences, further studies, possibly using both nuclear and mitochondria markers, would lead to better nodal supports for construction of a phylogenetic tree and provide insights on mechanisms of formation of latitudinal diversity gradients of pelagic copepods.

The larger OTU numbers in the mesopelagic than in epipelagic layers are consistent with previous studies that documented vertical diversity gradients in zooplankton [48, 49], and fine-scale segregations of vertical distributions that have been reported in the mesopelagic copepods [50, 51]. The high species richness but low phylogenetic diversity in mesopelagic layers indicated co-existence of many genetically similar species in the mesopelagic layer. Therefore, a recent rapid increase of species numbers during the evolutionary history is suggested in specific taxonomic groups (e.g., Augaptilidae and Scolecithrichidae), leading to fine-scale niche partitioning by closely related species in the relatively stable environment of the mesopelagic layer. The large vertical gradients in environmental factors such as temperature and chl-*a* may be associated with the larger variety of taxonomic groups found in the epipelagic layer, because vertical changes of environments are considered one important factor in maintaining planktonic taxa diversity [52]. In contrast with our results, in marine bacteria, more phylogenetically related OTUs are present in shallow water than in deep water [53]. This indicates that the mechanisms driving vertical diversity patterns may differ among taxa and that the recent rapid increase of species numbers in the mesopelagic layer may be unique in copepods. Although cold-water species are hypothesized to be ancestral to deep water species [42], this evolutionary process was not clearly shown in our metabarcoding study. The evolutionary processes responsible for the differences in the mesopelagic and epipelagic waters might be complex and different in each taxonomic group of oceanic copepods [45]. Copepods may have diversified over an evolutionary history, ultimately leading to the current co-existence of many species at low latitudes, especially in the mesopelagic layer where food sources are limited.

This metabarcoding study of copepods provided a first large-scale view of both horizontal and vertical changes in copepod community composition and diversity in the Pacific and Arctic Oceans, which have been formed through evolutionary processes and maintained in current environmental conditions. The results indicate that both the epipelagic and mesopelagic copepod communities may be vulnerable to anticipated global warming. This highlights the importance of understanding biogeography, community structure, and the mechanisms that create current patterns of biodiversity in order to monitor the effects of rapid environmental change. In addition to the distribution patterns of the dominant OTUs herein presented, all representative sequences and large-scale distributions of sequence reads for the OTUs produced in this study are available in the Dryad repository and can be easily included in a future study using the same method. Ultimately, more comprehensive sampling of zooplankton and the further development of molecular techniques will enable us to gain a deeper understanding of the mechanisms that drive diversity in copepods and other marine pelagic taxa.

## Materials and Methods

### Field sampling

Zooplankton samples were collected from 73 sampling stations across the Pacific and Arctic Oceans during ten research expeditions from 2011 to 2017 (Fig 1a). No specific permissions were required to conduct the samplings, as this study did not involve endangered or protected species. Samples in the Pacific Ocean were collected from the epipelagic (0–200 m), upper mesopelagic (200–500 m), and lower mesopelagic (500–1,000 m) layers using vertical tows of a vertical multiple plankton sampler (VMPS) with a 0.5 m^2^ mouth-opening area and 100-µm mesh. The samples in the Kuroshio regions were collected in three different seasons using a VMPS with a 0.25 m^2^ mouth-opening area and 100-µm mesh. For a subset of samples at a depth < 200 m in the Arctic Ocean, we used a North Pacific Standard Plankton (NORPAC) net, with a 100-µm mesh, to collect epipelagic samples using vertical tows from 5 m above the sea floor to the surface. A total of 205 bulk zooplankton samples were collected and immediately preserved in 99 % ethanol. The ethanol was replaced within 24 h of initial preservation, and samples were kept at 4 °C or −20 °C. Detailed information on the samples are listed in S1 Table. Vertical temperature and salinity profiles were obtained using a conductivity, temperature, and depth (CTD) system (SBE-911 plus, Sea-Bird Electronics). Dissolved oxygen content was measured using an SBE-43 dissolved oxygen sensor (Sea-Bird Electronics). Water samples were collected using Niskin bottles attached to the CTD system and were filtered using Whatman GF/F filters for chlorophyll-*a* (chl-*a*) analysis. Chlorophyll-*a* was extracted with *N*,*N-* dimethylformamide, and its concentration was analyzed using a Turner fluorometer [54]. Mixed layer depth (MLD) at each location was calculated by the depth at a temperature of ΔT = 0.2 °C from the temperature at 10 m depth [55].

### High-throughput sequencing and bioinformatic analysis

The metabarcoding analysis of copepods was mainly performed according to methods described in a previous paper [28]. Genomic DNA was extracted using the Gentra Puregene Cell and Tissue Kit (QIAGEN) from the 205 environmental zooplankton samples and two mock community samples containing morphologically identified copepod species (S1 Fig). The 19 environmental samples at epipelagic layer in the tropical and subtropical Pacific were previously analyzed using Roche 454 or Illumina MiSeq in our previous studies [26, 28], and the same DNA or raw sequence data were used in this study. The DNA concentration of each sample was measured with a Qubit 2.0 Fluorometer (Life Technologies). Library preparations were conducted using three-step polymerase chain reaction (PCR) using KOD Plus Version 2 (Toyobo) in a total of 25 μL reaction mixtures containing 13 μL distilled water, 2.5 μL 10×buffer, 2.5 μL dNTPs (2 mM), 1.5 μL MgSO_4_ (25 mM), 1.5 μL of each primer (5 μM), 0.5 μL KOD Plus polymerase, and 2 μL template DNA. The first PCR amplified the large ribosomal subunit (LSU) D2 region (approximately 400 bp) using template DNA (1 ng/μL) and a primer pair of LSU Cop-D2F (5ʹ-AGACCGATAGCAAACAAGTAC-3ʹ) and LSU Cop-D2R (5ʹ-GTCCGTGTTTCAAGACGG-3ʹ) [27]. The first PCR cycling included denaturation at 94 °C for 2 min, followed by 22 cycles of 10 s denaturation at 98 °C, 30 s annealing at 58 °C, and 1 min extension at 68 °C, with a final extension set at 68 °C for 7 min. PCR products from the first PCR cycle were diluted (1/20) with distilled water and used as template DNA for the second PCR, and then the process was repeated using the products the second PCR cycle template DNA for the third PCR cycle, which attached an adaptor and dual-index sequences for sequencing runs on Illumina MiSeq. PCR cycles were set at eight cycles with an annealing temperature of 50°C for the second PCR and 59 °C for the third PCR. Final PCR products were purified using a QIAquick PCR Purification Kit (QIAGEN), and the concentration of the purified PCR product was measured with a Qubit 2.0 Fluorometer. The quality of the final PCR products was confirmed by the Agilent DNA High Sensitivity Kit on the Bioanalyzer (Agilent). High-throughput sequencing runs were performed on the final PCR products using an Illumina MiSeq and 2 × 300 bp paired-end sequence reads were obtained. In this study, we firstly carried out bioinformatic analysis only using data of mock communities to determine parameters for OTU clustering. Data of mock communities were also analyzed together with environmental communities to validate accuracy of bioinformatic analysis of large sequence dataset. In bioinformatic procedure, raw paired-end reads were initially quality-filtered to remove sequences with < 100 bp length or average < 30 quality in every 30 bp using Trimmomatic [56]. Paired-end reads were merged in MOTHUR v.1.39.5 [57]. A quality-filtering step in MOTHUR removed sequence reads containing no ambiguous base and no primer sites. Quality-filtered sequence reads were classified into taxonomic groups using a naïve Bayesian classifier [58] with a threshold > 80 % to detect only copepod sequences. The reference dataset included manually curated 28S D2 sequences of 257 copepods and 36 other metazoan taxa. Further quality-filtering retained only copepod sequences containing ≤ 6 homopolymer and 300–420 bp length after removing primer sites. An equal number of sequence reads were selected in each sample, and all sequence reads were aligned using the add-fragments option in Multiple Alignment using Fast Fourier Transform (MAFFT) with the default setting [59]. Sequences that were not aligned with reference data were removed. We performed a single-linkage pre-clustering, removal of a singleton read, and chimera removal by UCHIME both with and without a reference dataset for the alignment sequences [60]. The final copepod sequence reads were subsampled again, and sequence differences were calculated among sequence reads without considering indels. OTU clustering was performed using OptiClust [61]. Following Hirai et al. [28] and the results of mock community analyses, we used a criterion of OTUs at the 98.5 % similarity threshold and a minimum sequence reads for OTUs of ≥ 8, to avoid overestimating diversity and to maintain an appropriate taxonomic resolution. The representative sequence of each copepod OTU was obtained based on the most abundant sequence reads. Detailed commands and datasets used in bioinformatic analysis are available in Dryad repository. All the following community and diversity analyses were conducted using OTUs, and taxonomy of each OTU was determined at the family level into the orders Calanoida and Cyclopoida (including Poecilostomatoida) and at the order level in other copepod groups.

### Copepod compositions based on presence/absence of OTUs

The broad-scale patterns of copepod compositions were investigated based on the presence or absence of OTUs. Cluster and multidimensional scaling (MDS) analyses were performed using Bray–Curtis similarity (Sørensen similarity in the case of presence/absence data). Clustered groups at 0–1,000 m depth were compared using Temperature-Salinity diagrams (T-S diagrams), which indicated the effects of water mass distributions on the community compositions of copepods. Permutational analysis of variance (PERMANOVA) was conducted to test the differences among clustered groups, which were determined based on community similarity and geographical regions. PERMANOVAs were conducted using Type III sums of squares and the unrestricted model. The variables explaining OTU composition were analyzed using a distance-based linear model permutation test (DistLM). The best models were selected using a step-wise selection procedure based on Akaike information criteria (AIC). The environmental factors included average water temperatures, salinity, and dissolved oxygen concentration, chl-*a* concentration, and MLD. The geographical factors were latitude and longitude. In the analysis of the epipelagic layer, we used average water temperature from 0 to 200 m, because of its higher correlation to community composition, rather than SST. In the analyses of mesopelagic layers and throughout the sampling layers (0–1,000 m), environmental valuables at the epipelagic layer were also included to evaluate the effects of the upper-layer environments. The community analyses were conducted in PRIMER version 7 with the PERMANOVA+ add-on [62, 63]. All permutation-based tests were conducted using 999 permutations.

### Quantitative analysis using sequence reads of OTUs

In addition to presence/absence of OTUs, quantitative data of sequence reads were analyzed for community structures of copepods and distribution patterns of major OTUs. Cluster analysis and PERMANOVA were conducted for all samples using the log_x+1_-transformed proportions of sequence reads in OTUs. The major OTUs were selected based on the results of a Similarity Percentage (SIMPER) analysis. The SIMPER analysis can select OTUs contributing to Bray–Curtis similarity within a group or dissimilarity between groups; thus, this analysis is useful for selecting major OTUs with distribution peak of sequence reads in a specific cluster group. The top five OTUs or OTUs contributing to at least 70 % of community similarity were selected as major OTUs in each cluster group. A BLAST search against the NCBI database was carried out for the representative sequences of major OTUs in order to obtain detailed taxonomic information. Cluster analysis was also performed for major OTUs based on their distribution patterns, and spatial distributions of sequence reads were compared among major OTUs.

### Spatial pattern of copepod diversity

The spatial patterns of the OTU numbers in each sample were compared at each layer and throughout the sampling layers. The evenness for each sample was evaluated using the Simpson diversity index. The phylogenetic diversity was calculated based on the average genetic distance between OTUs based on Kimura’s two-parameter nucleotide substitution model, which is a commonly used genetic distance for zooplankton including copepods [23]. Spatial differences in copepod diversity were evaluated using non-parametric Kruskal–Wallis tests and Dunn’s tests using SPSS 21.0 (IBM Corporation). The effect of environmental variables on OTU numbers were investigated using generalized linear models (GLMs) in R 3.5.0 [64]. We used the same environmental variables as in the DistLM analyses except for water temperature. SST was used as the temperature parameter in the epipelagic layer, because SST is more closely correlated to OTU numbers than average temperature. Based on the results of the dispersion tests, a negative binomial distribution with the log link function was used for the GLMs for each layer. The best model was selected based on AIC values obtained through backward selection of effective environmental variables. Based on the McFadden’s pseudo R^2^, the goodness of fit for the best model was estimated [65].

### Copepod biogeography and phylogenetic relationships

One of the advantages of metabarcoding analysis is the availability of representative sequences of OTUs. The sequences were used to investigate effects of evolutionary processes on large-scale biogeographic patterns of copepods. An evolutionary process related to latitudinal diversity gradients was particularly focused in the phylogenetic analysis. The biogeographic pattern of each OTU was determined based on distribution of sequence reads. If the highest proportion of sequence reads was observed in the arctic or subarctic, this OTU was classified as an OTU with a cold-water distribution. We analyzed OTUs with ≥ 15 sequence reads in a single sample. The proportions of distribution patterns were investigated in each taxonomic group. The taxonomic treatment of groups of copepods followed that used in Blanco-Bercial et al. [30] and Khodami et al. [66]. Phylogenetic analyses were conducted in the taxonomic groups containing OTUs with both high-latitude (cold water) and low-latitude distributions (warm water). We added outgroups to each taxonomic group, and sequences were aligned again using MUSCLE [67]. After selection of a best-fit nucleotide substitution model, maximum likelihood analyses were performed with 100 bootstrap replicates to assess nodal support using MEGA 6.0 [68]. The nearest genetic distances of the OTUs were also calculated using Kimura’s two-parameter nucleotide substitution model, and average values were compared between OTUs with cold and warm distributions.

## Supporting information

**S1 Fig. Comparison of operational taxonomic units (OTUs) and reference sequences in mock community analyses.** Scale bar indicates genetic distance (p-distance). Note that bioinformatics of mock community analyses were performed together with all environmental community data for validation accuracy of data analyses.

**S1 Table. Metadata of the environmental zooplankton samples used in this study.**

**S2 Table. Summary of the preliminary analyses of mock community.** These preliminary analyses of mock communities were performed to determine similarity threshold to cluster OTUs and abundance threshold to remove rare and erroneous OTUs. Estimated numbers of OTUs are based on reference sequences obtained by Sanger sequencing. An abundance threshold of 8 and a similarity threshold of 98.5%, which are values for environmental community analysis, were used for comparing different values of similarity and abundance threshold, respectively. Target OTUs are numbers of OTUs identified as copepod species contained in a mock community, and non-target OTUs are other copepod OTUs without high similarity to target species. Note that seven species in mock community 2 were incubated after sampling to remove gut contents, but 33 species without incubation were used for mock community 1. The analyses were based on the preliminary analyses only using data of mock communities.

**S3 Table. Summary of the permutational analysis of variance (PERMANOVA).** Copepod community compositions (presence/absence of Operational Taxonomic Units) were compared at each sampling layer between cold-water and warm-water groups and throughout the water column. The effect of sampling layers (epipelagic and mesopelagic) on community composition was investigated for all sampling locations and for warm-water regions. The effect of cluster group on copepod community compositions was investigated for each sampling layer and throughout the water column. The differences among cluster groups based on quantitative data of the sequence reads were also analyzed.

**S4 Table. Summary of the distance-based linear model permutation test (DistLM) for copepod community based on sequence reads.** The best model of environmental variables explaining copepod community was selected based on Akaike information criteria (AICs) for all locations in each sampling layer [epipelagic (0–200 m), upper mesopelagic (200–500 m), and lower mesopelagic (500–1,000 m)]. Pseudo–*F*, *P*–value, and explained variation attributable to the model are indicated for each environmental variable.

**S5 Table. Summary of Kruskal–Wallis and Dunn’s test results for copepod diversity.** Operational Taxonomic Unit (OTU) numbers, Simpson index, and phylogenetic diversity were compared among areas at each sampling layer [shallow: epipelagic (0–200 m), middle: upper mesopelagic (200–500 m) and deep: lower mesopelagic (500–1,000 m)] and among each sampling layer at each area (see Fig. 7). Where Kruskal–Wallis tests were significant, groups with adjusted *P* < 0.05 in pairwise comparisons using Dunn’s tests are listed. Ar = Arctic, Sa = subarctic, Tra = Transition, Ns = North subtropical gyre, Tro = Tropical, Ss = South subtropical.

## Acknowledgments

We thank the captains and crews of the RV Hakuho–Maru, RV Shinsei–Maru, and FRV Soyo–Maru for their assistance with field collection. We also thank the three anonymous reviewers for suggestions on the manuscript. Sequencing runs on the Illumina MiSeq were carried out in FASMAC Co., Ltd.

## Financial Disclosure

This work was supported by grants from the Japan Society for the Promotion of Science (grant nos. 247024 to J.H. and 24121004 to A.T.). The funder had no role in study design, data collection and analysis, decision to publish, or preparation of the manuscript.

## Data Availability

Raw sequence data are available in the NCBI/EBI/DDBJ Sequence Read Archive (BioProject accession PRJDB7448). Detailed commands and datasets used in bioinformatic analysis are available in Dryad repository (doi:10.5061/dryad.x95x69pdt). All lists of OTUs and sequence reads in each sample are also available with representative sequences in the same Dryad repository.

## Author Contributions

**Conceptualization:** Junya Hirai, Aiko Tachibana, Atsushi Tsuda.

**Formal Analysis:** Junya Hirai.

**Funding Acquisition:** Junya Hirai, Atsushi Tsuda.

**Investigation:** Junya Hirai, Aiko Tachibana, Atsushi Tsuda.

**Methodology:** Junya Hirai.

**Project Administration:** Junya Hirai.

**Resources:** Junya Hirai, Aiko Tachibana, Atsushi Tsuda.

**Supervision:** Atsushi Tsuda.

**Visualization:** Junya Hirai.

**Writing – Original Draft Preparation:** Junya Hirai.

**Writing – Review & Editing:** Junya Hirai, Aiko Tachibana, Atsushi Tsuda.

